# A synthetic method to assay polycystin channel biophysics

**DOI:** 10.1101/2024.05.06.592666

**Authors:** Megan Larmore, Orhi Esarte Palomero, Neha P. Kamat, Paul G. DeCaen

## Abstract

Ion channels are biological transistors that control ionic flux across cell membranes to regulate electrical transmission and signal transduction. They are found in all biological membranes and their conductive state kinetics are frequently disrupted in human diseases. Organelle ion channels are among the most resistant to functional and pharmacological interrogation. Traditional channel protein reconstitution methods rely upon exogenous expression and/or purification from endogenous cellular sources which are frequently contaminated by resident ionophores. Here we describe a fully synthetic method to assay functional properties of polycystin channels that natively traffic to primary cilia and endoplasmic reticulum organelles. Using this method, we characterize their oligomeric assembly, membrane integration, orientation and conductance while comparing these results to their endogenous channel properties. Outcomes define a novel synthetic approach that can be applied broadly to investigate channels resistant to biophysical analysis and pharmacological characterization.

## INTRODUCTION

Ion channels are pore forming integral transmembrane proteins essential for all cellular lifeforms^1, 2^. At the plasma membrane, ion channels are responsible for generating long range bioelectric conduction in excitable eukaryotic cells (e.g. neurons), and within prokaryotic colonies (e.g. bacteria biofilms), and filamentous colonies of archaea^3-5^. Here, they contribute to fundamental vital cell processes including division, signal transduction, and ionic homeostasis^1^. In eukaryotic organelle membranes, ion channels are integral to a wide range of functions including energy production (mitochondria), and the maintenance of defining compartmental conditions such as Ca^2+^ gradients (endoplasmic reticulum), pH (lysosome) and redox states (peroxisome)^6-8^. The conductive states of channels are precisely controlled by their molecular conformations which are unique among their phylogenetic families and subfamilies^9^. More than 400 human genes encode ion channel subunits, many of which are altered by variants that impact organ function and development^10-12^. These so called ‘Channelopathies’ manifest in various human diseases such as cardiac arrhythmias, neurological conditions, and cystic kidney diseases^13, 14^. Besides their association with disease-causing variants, channels are important therapeutic targets for treating various pathophysiologies and represent the second largest target among the existing FDA-approved drug portfolio^15-17^. However, many members of the so called “dark genome” of understudied proteins encode putative ion channels that are implicated in human disease but remain resistant to functional characterization^18^. Furthermore, channels which localize to cellular compartments in low quantities present a significant challenge to assay for drug screening purposes^19^. Thus, these observations warrant the present investigation of a novel methodological approach to characterize ion channel biophysics and pharmacology.

Voltage clamp electrophysiology incarnated in either planar or glass electrode designs provide direct, real-time and the highest available fidelity measurements of the ion channel conductive states. In the on-cell or inside-out configurations, their opening and closing conformational changes (i.e. gating) are captured as stochastic single channel currents after forming high resistance seals (>10 MΩ) at the interface of the electrode against a biological membrane^20^. This conventional electrophysiology technique is commonly used to characterize the properties of plasma membrane channels and typically involves either recording from an endogenous cell source or from a cell line expressing the channel transgene ^21, 22^. However, capturing the biophysical properties of organelle channels and those from bacteria can be mired by electrode inaccessibility to inner membranes. As a work around, investigators have employed various channel reconstitution methods which typically consist of several steps^23, 24^. First, the channel of interest is expressed and immunopurified from a biological cell source, then it is reconstituted into a synthetic or into biologically derived lipid bilayers or vesicles. While there are many examples where heterologous approaches have faithfully reproduced the functional properties of channels from native sources, these preparations are frequently contaminated by endogenous ionophores from the host cell lines, even after purification^25, 26^.

To address this, we have developed a completely synthetic method to assay ion channel biophysics. The approach combines advances in cell-free protein expression (CFE) and reconstitution of the synthetic channel protein into giant unilamellar vesicles (GUV) where their single channel properties can be measured using voltage-clamp electrophysiology^27-29^. CFE is a form of *in vitro* protein synthesis, utilizing purified cellular machinery (ribosomes, tRNAs, enzymes, cofactors, etc.) needed to directly transcribe and translate user-supplied DNA plasmid encoding an ion channel. The unmodified channel proteins are reconstituted into GUVs— cell-sized model membrane systems derived from electrolysis of synthetic lipid mixtures. To validate this method, we characterized PKD2 and PKD2L1, two members of the polycystin subfamily of transient receptor potential (TRP) ion channels. PKD2 and PKD2L1 are highly homologous subunits with six transmembrane segments and shared overall protein folding when structurally assembled as homotetrameric channels^30-35^. Polycystin subunits can also form heteromeric ion channel complexes with several members of the TRP family and traffic to the primary cilia and ER organelle membranes^36^. Both features present challenges for experimentalists to functionally characterize these unique channel properties within their endogenous cell membranes. The medical and biological importance of polycystins is underscored by PKD2 variants associated with autosomal dominant polycystic kidney disease (ADPKD), and this channel’s role in fertility and conferring right-left symmetry in embryonic development^37-39^. While PKD2L1 variants have yet to be linked to human disease, its loss of expression results in epilepsy susceptibility and autism -like features in mice^40, 41^. In this report, we stepwise express and confirm protein expression of polycystin channels using the CFE method; reconstitute channel protein in GUVs containing distinct lipid components; assess correct membrane orientation using self-labeling saturated N-heterocyclic building blocks (SNAP) protein chemistry and evaluate channel properties using electrophysiology^42^. Outcomes define a novel reductionist and generalizable approach to study ion channels resistant to biophysical characterization.

## RESULTS AND DISCUSSION

We began by carrying out CFE in vitro synthesis of PKD2L1 protein in the presence and absence of lipid vesicles. Plasmid DNA encoding human PKD2L1 with C-terminally tagged green fluorescent protein (PKD2L1-GFP) was added to the cell-free expression components (PURExpress, New England Biolabs) and induced protein translation by heat (**Figure 1A**, See methods). Each reaction produced 8 ± 3.5 ng/μl of synthetic channel protein as estimated by a standardized GFP absorbance assay (**Figure 1— figure supplement 1A**). The synthetically derived channel protein identity was confirmed using two methods. First, by western blot of the cell-free reaction where the PKD2L1-GFP protein was SDS-PAGE gel separated from the reactants and detected by an anti-GFP monoclonal antibody (**Figure 1B**). Second, the PKD2L1 protein was confirmed by mass spectroscopy with 46% coverage (**Figure 1— figure supplement 1B**,**C**). Since polycystins are transmembrane proteins, we hypothesize that channel membrane incorporation would be facilitated by including lipid substrates into the CFE reaction. Thus, we compared channel synthesis in the presence or absence of small unilamellar vesicles (SUVs) comprised of 1,2-diphytanoyl-sn-glycero-phosphocholine (DPhPC; 4ME16:0PC) and cholesterol (**Figure 1A**). We monitored PKD2L1-GFP protein production over time through fluorescence which is dependent on complete channel translation and correct GFP folding^43^. We observed a dramatic increase in fluorescence output which plateaued after 3 hours when CFEs reactions were supplemented with SUVs—and effect not observed in water-supplement (H_2_O) control reactions (**Figure 1C**)^44^. Minimal changes in fluorescence were detected when a control plasmid (Ctrl) encoding a non-fluorescent protein (dihydrofolate reductase) was used in the reaction. Polycystin channels function as tetramers, thus we examined CFE derived PKD2L1 oligomeric assembly in SUVs using fluorescence-detection size-exclusion chromatography (FSEC). As controls, we tested recombinant (cell-derived) GFP and GFP tagged polycystin proteins which produced monodispersed peaks in the fluorescent signal which provided a reference for their respective elution times off the column (**Figure 1— figure supplement 1D**). Although not as robust, the fluorescence signal from CFE+SUV derived PKD2L1 protein produced a symmetrical peak at the expected elution time of channel tetramers, along with fractions which may correspond to unassembled protomers (**Figure 1—figure supplement 1E**). Taken together, these results demonstrate the feasibility of synthesizing full length polycystin channels using the cell free expression system, and the enhancement of transmembrane protein synthesis and channel assembly through lipid vesicle incorporation during these reactions.

**Figure 1.**
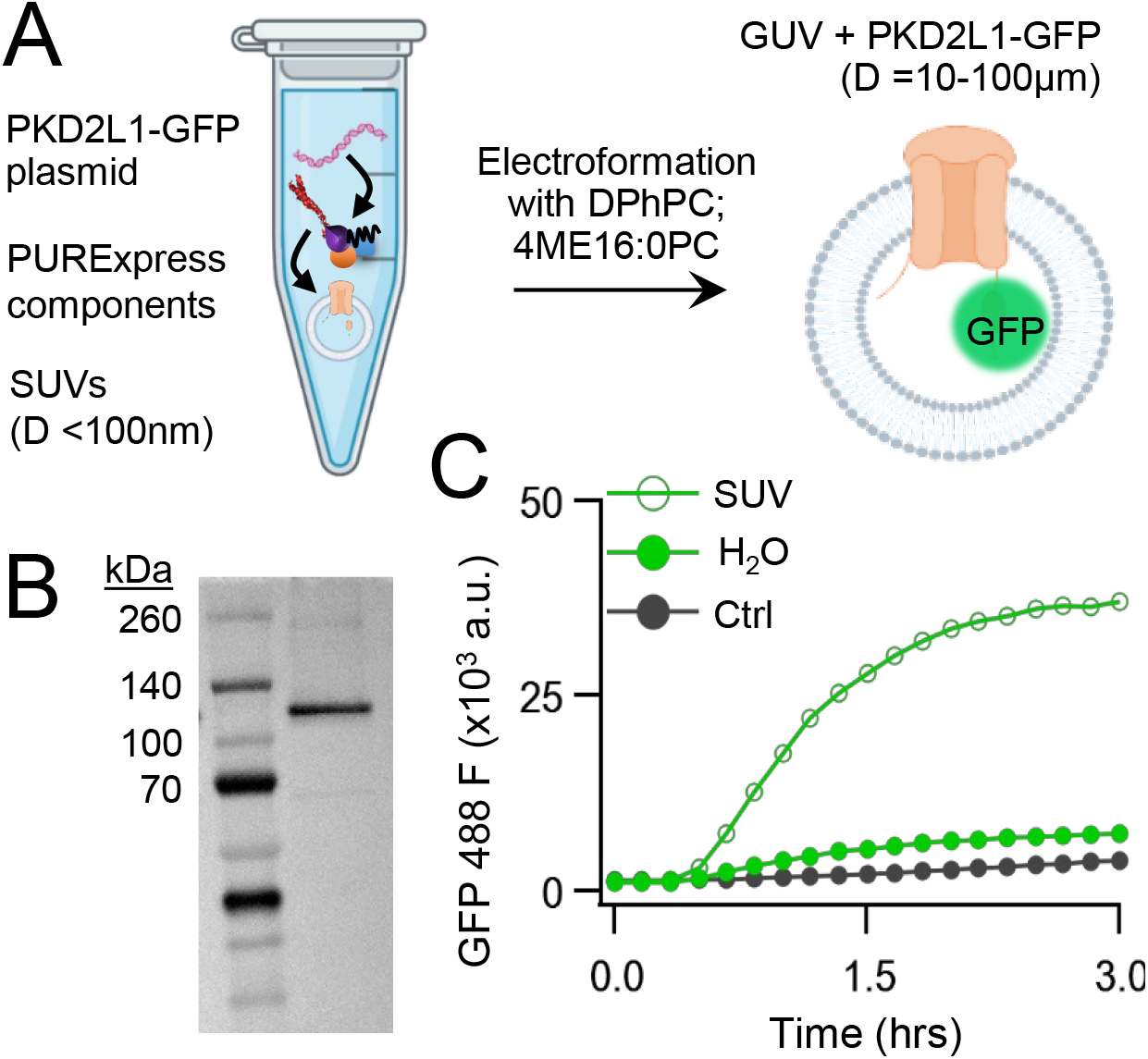
Cell-free expression of synthetic PKD2L1 protein and incorporation into lipid vesicles. **A**) Schematic of cell-free protein expression into synthetic lipid vesicles and subsequent electroformation with Vesicle Prep Pro (Nanion). **B)** Full length PKD2L1-GFP protein detected by Western Blot after cell-free expression into vesicles. **C**) Monitored fluorescence over time of cell-free expressed PKD2L1-GFP and a non-fluorescent control plasmid produced in the presence or absence of lipid vesicles.

One caveat of membrane protein vesicle reconstitution is the potential for misorientation after integration. To assay channel orientation, we synthesized PKD2L1 with a C-terminal SNAP-tag fusion protein (PKD2L1-SNAP) in SUVs then electroformed them into GUVs for the assay. The SNAP-tag specifically reacts with fluorescent O2-benzylcytosine derivatives and, depending on the derivatives membrane permeability, will react with lipid integrated proteins based on the tag’s accessibility (**Figure 2— figure supplement 1**). GUVs containing CFE synthesized PKD2L1-SNAP were formed by electroformation after passing current through indium tin oxide slides treated with dried SUV-channel protein mixture^45^. We then added two SNAP fluorescent derivatives, membrane permeable SNAP-Cell Oregon Green (Cell488) and membrane impermeable SNAP-Surface Alexa Fluor 647 (Surface647), to monitor orientation-dependent membrane protein reactivity with the SNAP label (**Figure 2A, Figure 2— figure supplement 1**). We hypothesized there will be two fluorescent outcomes. First, if all channels orient correctly, then we should see only Cell488 fluorescence with no Surface647 at the membrane (**Figure 2A**). Second, if the channels are in opposite or in mixed orientations, then we expect to see dual fluorescence of Cell488 and Surface647 (**Figure 2B**). We imaged over 60 GUVs and found 38.5% of the vesicles exhibited sole Cell488 fluorescence— indicating their correct channel orientation within this population (**Figure 2B, C**). Importantly, none of the vesicle membranes labeled with both Cell488 and Surface647 while retaining a clear lumen— suggesting that the population of GUVs containing misoriented channels was nominal. In some cases, vesicles can encapsulate dye through compromised integrity or mechanisms other than membrane permeability. This is apparent when fluorescence can be seen in the vesicle lumen, rather than on the membrane (**Figure 2C**). While the encapsulated fluorescent vesicles account for most of the vesicles imaged, there were no vesicles seen with Cell488 and Surface647 fluorescence at the membrane with a clear lumen (**Figure 2B, C**). Based on these results, we conclude that our cell-free synthesized PKD2L1 channels are successfully reconstituted in GUVs in the correct orientation, and this preparation is suitable to assay PKD2L1 channels using electrophysiology.

**Figure 2.**
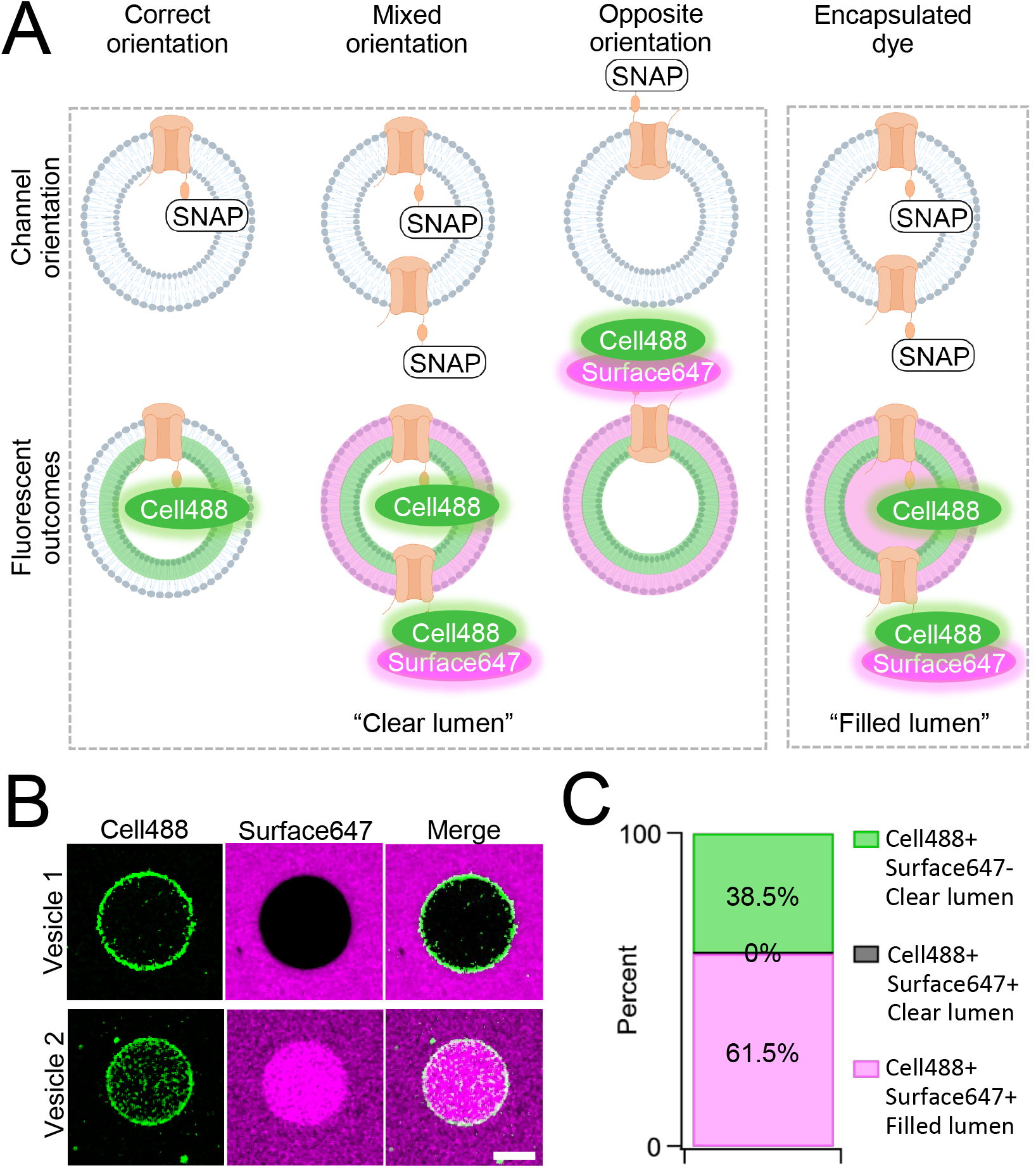
Orientation of synthetic PKD2L1 channels in vesicles. **A**) Schematic of possible ion channel orientation outcomes from PKD2L1 cell-free expression (top) and hypothesized fluorescence results when Cell488 and Surface647 added (bottom). **B**) Fluorescent confocal images from the SNAP-tagged vesicles. The scale bar represents 10 μm for all images. **C**) Vesicle percentage depicts the percent of vesicles with each fluorescent output, (N=65 vesicles).

Native and transgene expressed PKD2L1 channels conduct monovalent cations, thus we established symmetric potassium ion (K^+^) recording conditions to measure synthetic polycystin currents from GUVs. GUVs containing PKD2L1 were voltage-clamped using 4-7 MΩ resistance (R) glass electrodes in the inside-out membrane patch configuration (**Figure 3A**). Single channel events were frequently observed while establishing high resistance seals (R >10 GΩ), however the majority of PKD2L1 GUV patches were unstable during voltage steps and these results were excluded from the final analysis. We hypothesize that patch instability and low seal resistance likely results from over-incorporation of PKD2L1 tetramers into the GUV membranes, as supported by the FSEC data. In addition, membrane instability was not observed from empty GUV recordings, suggesting that opening of incorporated CFE synthesized polycystins is likely responsible for patch destabilization. From the stable recordings, two magnitudes of single channels were readily observed from GUVs containing PKD2L1, suggesting unique full and sub-conductance states (**Figure 3B, C, Table 1**). In some recordings, only one conductance predominates, which can be estimated from recordings from individual GUVs (**Figure 3— figure supplement 1A, B**). While in other recordings, both full (*FC*) and sub conductive (*SC*) states can be identified by histogram analysis of the unitary current (**Figure 3— figure supplement 1C, D**). Importantly, no single channel events were observed from GUVs (N= 11) derived from CFE reactions with the empty plasmid— indicating that the measured conductance is not due to contaminates from lipid or cell-free reagents (**Figure 3B**). To assess the selectivity of the synthetic PKD2L1 pore, we substituted the pipette K^+^ charge carrier for methyl-D-glucamine ions (NMDG^+^). We did not observe any outward single channel currents (i.e. towards the bath), indicating the large cation was not permeable through PKD2L1 which is consistent with previous reports (**Figure 3D, E**)^46, 47^. To determine the feasibility of using this approach to assess the function of other polycystin channels, we followed the same steps to assay PKD2 channel biophysics. As observed in our PKD2L1 results, unitary single channel currents of synthetic PKD2 channels reconstituted in GUVs yield sub- and full conductances, which were terminated by substitution of K^+^ with NMDG^+^ in the electrode (**Figure 4A-C**; **Figure 3— figure supplement 1E-H**). To compare the properties of the synthetic and biologically derived channels, we recorded native PKD2L1 and PKD2 channel single channel currents from the primary cilia membranes of hippocampal neurons and inner medullary collecting duct (IMCD) cell line, respectively (**Figure 4—figure supplement 1**)^40, 48, 49^. Like the synthetic PKD2 and PKD2L1 channels, native polycystins produced sub and full K^+^ conductances with inward currents having the greater magnitudes. Here, the synthetic PKD2L1 GUV conductance approximates the native full and subconductances recorded from hippocampal primary cilia membranes cultured from neonatal mice (ARL13B-EGFP^tg^). However, the PKD2 K^+^ conductance magnitudes recorded from IMCD cilia were significantly smaller than those assayed using the CFE-GUV synthetic system (**Table 1**). These differences might arise from the lack of post-translational modifications (e.g. phosphorylation and N-glycosylation) to the synthetic PKD2 peptides, which are normally found in biologically derived channels ^35, 50, 51^. In addition, the GUVs are comprised of synthetic lipids which does not reflect the composition of organelle (cilia or ER) membranes of the cell^52^. Thus, while retaining the native ion selectivity and ion channel functionality despite their cell-free origin, synthetic PKD2 has different conductance magnitudes compared to cell derived channels, which presents a limitation of using this approach to recapitulating physiological channel functions.

**Figure 3.**
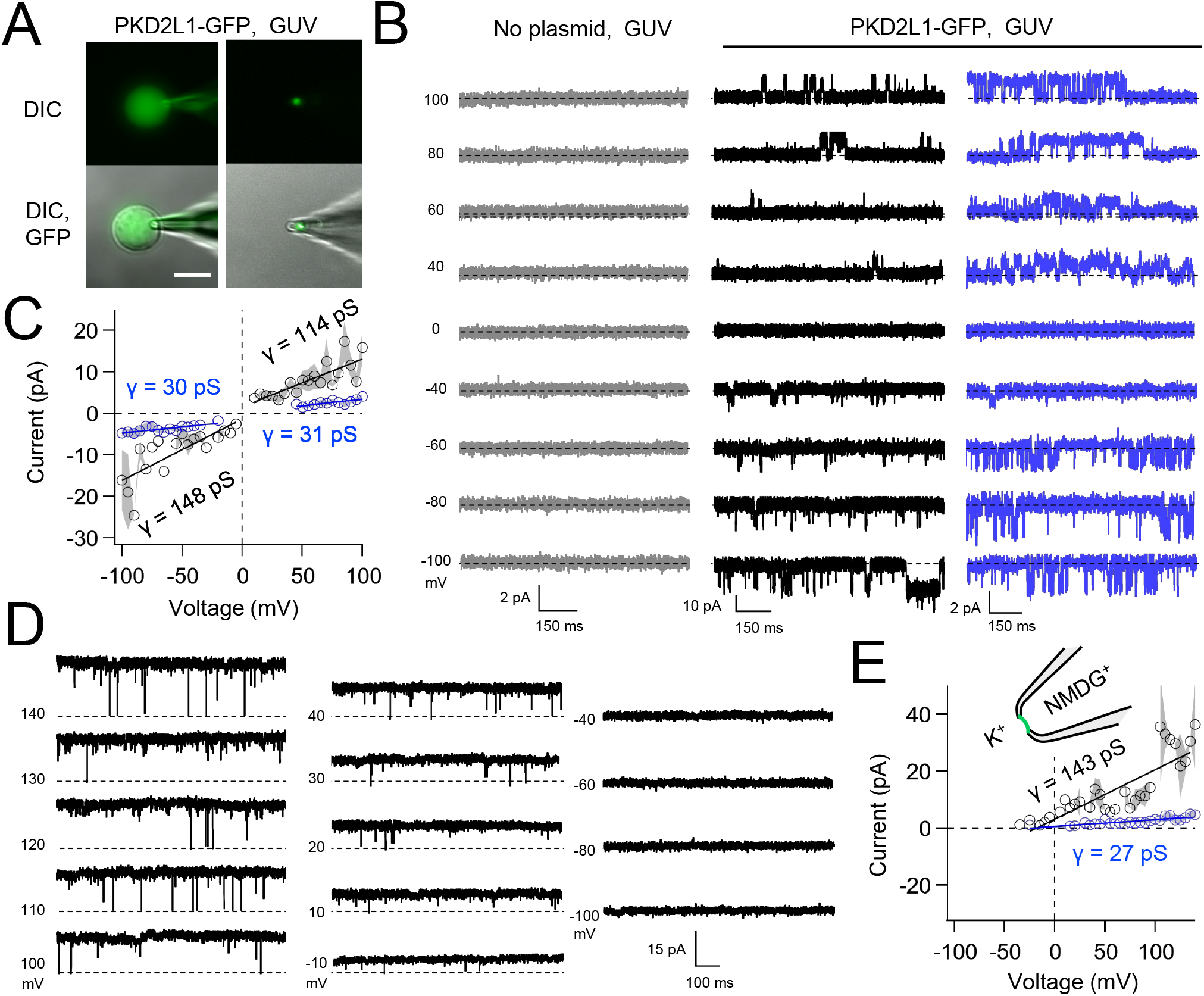
Synthetic PKD2L1 channels are functional in artificial membranes. **A**) Images of voltage clamped GUVs with incorporated PKD2L1-GFP channels. *Left*, establishing high-resistance seals in the on-cell patch configuration. *Right*, transitioning to the inside-our patch configuration. Scale bar = 20 μm. **B**) Example unitary single channel current records from GUVs reconstituted with or without PKD2L1 protein. Vesicles were patched using the symmetrical 150 mM K^+^ conditions (see methods) and PKD2L1 single channel current producing full and sub-conductances are colored black and blue, respectively. **C**) Average single-channel current amplitudes. Conductance (γ) estimated by fitting the average single channel currents to a linear equation. Error (grey) indicates SEM from N=3-8 GUVs. Several replicates lacked single channel openings at all test potentials. Conductance estimates from individual GUVs are shown in Figure 3—Supplement Figure 1A. **D**) PKD2L1 single channel current recorded using asymmetric cationic solutions, with 150 mM K^+^ in the bath and 150 mM NMDG^+^ in the pipette. **E**) Resulting average single-channel current amplitudes where no inward single channel current was detected (N=3-4 GUVs).

**Table 1.** Conductance properties of polycystins measured from GUV and cilia membranes.

| Polycystin channel and membrane context. | Major K<sup>+</sup> γ (pS) ± S.D. |  | Minor K<sup>+</sup> γ (pS) ± S.D. |  |
| --- | --- | --- | --- | --- |
|  | *inward* | *outward* | *inward* | *outward* |
| PKD2 (GUV membrane, cell free expression) | 282 ± 38 | 153 ± 32 | 23 ± 4 | 21 ± 3 |
| PKD2 (primary cilia membrane, endogenous murine IMCD cell line) | 144 ± 9 | 110 ± 6 | 46 ± 4 | 34 ± 4 |
| PKD2L1 (GUV membrane, cell free expression) | 148 ± 29 | 114 ± 28 | 30 ± 6 | 31 ± 7 |
| PKD2L1 (primary cilia membrane, endogenous hippocampal neurons) | 165 ± 10 | 113 ± 6 | 40 ± 4 | 29 ± 3 |

**Figure 4.**
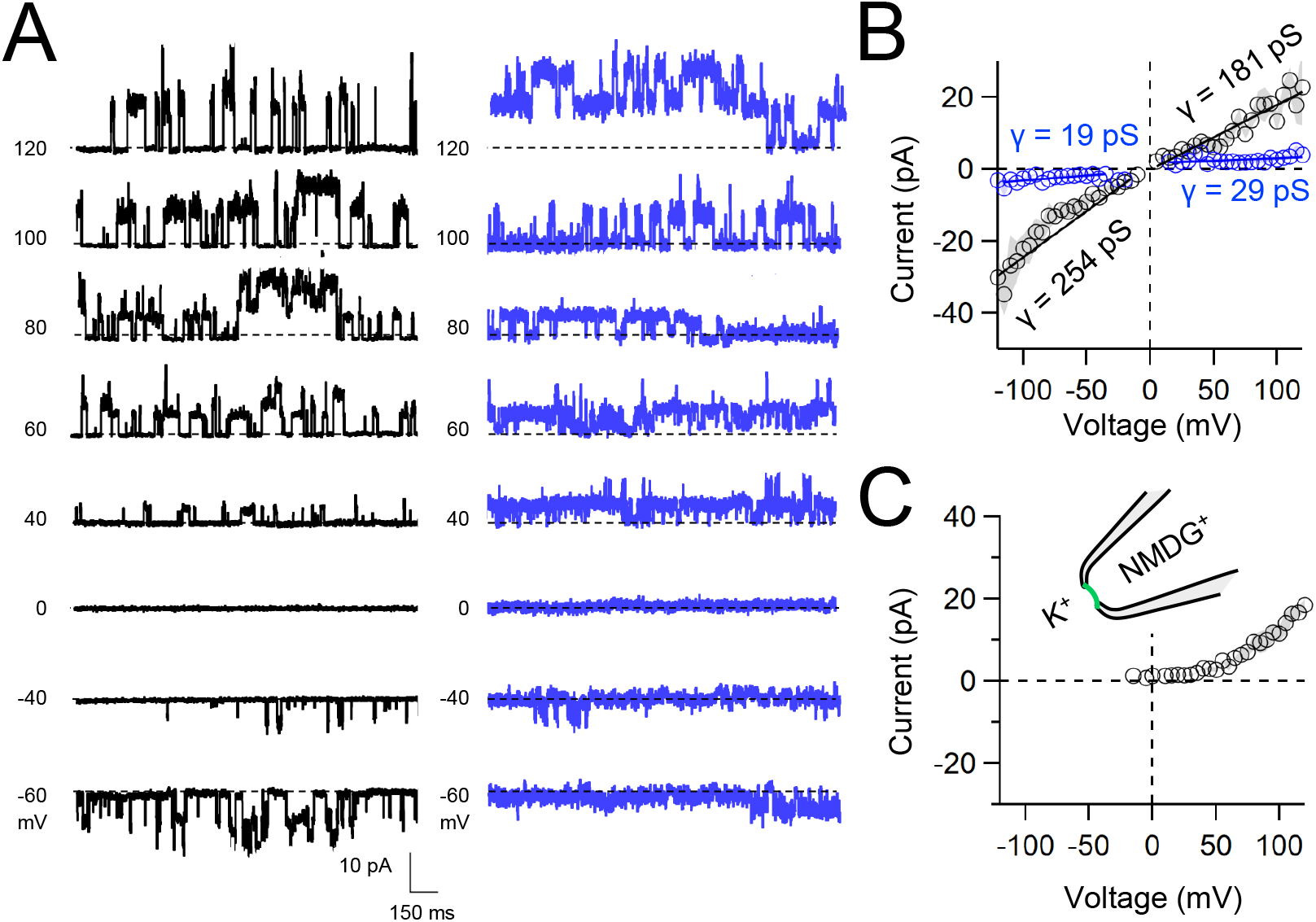
Functional synthetic PKD2 channels in artificial membranes. **A)** Example unitary single channel current records from GUVs reconstituted with PKD2 protein in symmetrical 150 mM K^+^ conditions producing full (black traces) and sub-conductances (blue traces), respectively. **B, C**) Average single-channel current amplitudes recorded using K^+^ or NMDG^+^ in the recording electrode solution. Conductance (γ) estimated by fitting the average single channel currents to a linear equation. Error (grey) indicate SEM from K^+^ (N=3-12 GUVs) and NMDG^+^ (N=2-5 GUVs) conditions.

In summary, we have established a synthetic approach to assay ion channel biophysics using two polycystin members to validate our method. Previously CFE has been used to study membrane protein integration, drug delivery, and the study of actin dynamics^29, 53, 54^. The novelty of our approach rests with the adaptation of CFE derived channels and their GUV reconstitution to carry out single channel electrophysiology experiments. The described method represents a highly reductionist approach to assay channel function which can be generalized to other channels resistant to characterization using traditional electrophysiology approaches. Furthermore, the CFE-GUV electrophysiology method can be leveraged for future inquiry into lipid-channel regulation and effects of channel subunit composition. PKD2-related protomers form heteromeric complexes (e.g. PKD1-PKD2; PKD1L1-PKD2L1) with PKD1-related polycystins which are notoriously difficult to assay using traditional electrophysiology techniques (as reviewed)^36^. Using the CFE method to co-synthesize and GUV reconstitute these subunits presents a potential avenue to assay their function with patch clamp recordings. The approach may be further developed into high throughput drug screening assays using cation reporters (e.g. Fura derivatives) or multi-well planer electrophysiology instruments^55, 56^. With our pipette patch electrodes, we observed considerable instability of our high resistant seals when the bath solutions were exchanged. Thus, future work could explore alternative membrane compositions (e.g. additional cholesterol) and stabilizing cationic conditions (internal CsF) to mitigate this effect. The folding and membrane integration of many ion channels require their association with stabilizing chaperone proteins produced in cells^57-59^. Thus, while the co-expression of chaperones with these channels using the CFS+GUV system is likely required for their functionality, the approach may also be leveraged to study chaperone-assisted folding *in vitro*.

## METHODS

### Protein production

Cell-free protein production was performed with PURExpress In Vitro Protein Synthesis Kit from New England Biolabs, Inc. (Ipswich, MA, USA). Both PKD2 and PKD2L1 are in a pET19b plasmid under T7 promoter. We utilized the manufacturer’s protocol with 1 mg target DNA and a maximum reaction volume of 30 μl. When appropriate, we added substituted DiH_2_O for SUVs. The reactions were placed in 37°C water bath or heated plate reader between 2 – 3 hours and then placed at 4°C for storage.

### Vesicle formation

Lipids 1,2-diphytanoyl-sn-glycero-phosphocholine (DPhPC 4ME16:0PC) and cholesterol (ovine) were obtained from Avanti Polar Lipids (Alabaster, AL, USA) and mixed in chloroform to the desired mol percentage, 95% DPhPC and 5% cholesterol. SUVs were formed following the previously described [25]. Briefly, lipids were reconstituted in chloroform in a glass vial and the chloroform was evaporated until a thin lipid layer is deposited on the bottom of the glass vial. The lipid layer is then placed under vacuum, -23 inhg, for >4 hours. Lipids are then rehydrated in 1ml of diH2O overnight at 60°C. The following day, lipids are vortexed and then passed through a 100 nm polycarbonate filter with the Mini-Extruder (Avanti Polar Lipids, Alabaster, AL USA) 7 times and stored at 4°C for two weeks. SUVs with or without channel incorporated are dried onto indium tin oxide coated glass slides from Nanion Technologies (Munich, Germany). The dried slides are placed on the Vesicle Prep Pro (Nanion Technologies) with a rubber o-ring and 300 mM sucrose. GUVs are formed using the standard program. GUVs are electroformed and used for electrophysiology experiments the same day. SUVs with channel incorporated are stored at 4°C for 3 days.

### Monitoring fluorescence and cell free protein synthesis quantification

We monitored fluorescent folding with PKD2L1 C-terminally tagged GFP during PURExpress reaction in the presence and absence of SUVs. GFP fluorescence was monitored every 10 minutes for three hours at 37°C on the BioTek Cytation5 Imaging Reader (Agilent, Santa Clara, CA, USA). Control plasmid was the PURExpress Control DHFR Plasmid (NEB, Ipswich, MA USA) with no fluorescent tag. Green fluorescent protein (GFP) standard curve was created from dilutions of Aequorea Victoria GFP His-tag recombinant protein (ThermoFisher Scientific, Cat. No. A42613) and measured on the BioTek Cytation5 Imaging Reader. A linear regression (Igor Pro, WaveMetrics, Portland, Or.) was used to create a standard curve. Target protein fluorescent measurements were made after in vitro protein synthesis for three hours.

### Western blotting

Western blotting was performed on PKD2L1–GFP plasmid after PURExpress protein expression in the presence of SUVs. SUV and protein mixture were separated on SDS-PAGE gel, Novex Tris-Glycine mini protein gels, 4-20%, 1.0 mm, WedgeWell format, (ThermoFisher, Waltham, Ma USA). The SDS-PAGE was run with 10 μl Spectra Multicolor Broad Range Protein Ladder (ThermoFisher Scientific, Cat. No. 26634). The gel was then transferred to Amersham Hybond P 0.45 PVDF blotting membrane (Cytiva, Cat. No. 10600029) and PKD2L1-GFP was detected with an anti eGFP monoclonal antibody (F56-6A1.2.3) (Invitrogen, Cat. No. MA1-952) diluted 1:1000 in TBS with 0.1% (v/v) Tween-20 and 5% (5/v) milk overnight at 4°C. The goat anti-mouse AF555 secondary (Invitrogen, Cat. No. A32727) diluted 1:5000 in TBS with 0.1% Tween-20 and 5% milk was incubated for 1 hour at room temperature.

### SNAP staining

Channel orientation was visualized with N-terminally tagged PKD2L1 with SNAP Tag sequence, (NEB, Ipswich, MA USA). After PKD2L1-SNAP-tag production and incorporation into GUVs, two SPAP-tag substrates were added to the solution, cell permeable SNAP-Cell Oregon Green and the cell impermeable SNAP-Surface Alexa Fluor 647, according to manufacturer’s instructions. Images were collected on Nikon A1 confocal microscope and all images were analyzed with Nikon Elements (Melville, N USA). Briefly, regions of interests were manually outlined around the vesicle membrane. Then Pearson’s correlation coefficients were measured for fluorescence overlap of the two substrates.

### Fluorescence-detection size-exclusion chromatography

As controls for the polycystin FSEC elution time, Aequorea Victoria GFP His-tag recombinant protein (ThermoFisher Scientific, Cat. No. A42613) and polycystin channel protein was harvested from lysates of 0.5-×10^9^ HEK293T (ATCC, Cat No. CRL-3216) PKD2^Null^ cell lines^60^ stably expressing PKD2-GFP and PKD2L1-GFP. Sythetic channel protein from 3 CFE reactions were synthesized in the presence of SUVs, as previously described. The SUVs were lysed using Dodecyl β-D-maltoside (DDM) (GoldBio) and protein supinates collected after centrifugation for 20 min at 20,000 rpm. Samples were then diluted in 50-100 μl of SEC running buffer containing (ml) 150 mM NaCl, 25mM HEPES, 1mM CaCl_2_-6H_2_O, 0.05% DDM, 0.005% cholesteryl hemisuccinate, pH 7. 50 μl of each sample were separated on an analytical size-exclusion column (Superose 6 5/150 GL; GE Healthcare) at 0.2 mL/min flow rate. Fluorescent proteins were detected (excitation: 480 nm, emission: 512 nm) using an RF-20Axs detector (Shimadzu, Japan).

### Isolation of primary hippocampal neurons and IMCD cell culture for cilia electrophysiology

All mice utilized in these procedures are housed in our AAALAC-approved Center for Comparative Medicine (CCM) at Northwestern University, Feinberg School of Medicine. NU Institutional Animal Care and Use Committee has approved this facility and was monitored by an Animal Care Supervisor as well as a veterinarian from the Division of Laboratory Animal Medicine (DLAM). All of those who handled animals and perform the approved protocols were properly trained prior to start of work to ensure no animal discomfort. Mice were anesthetized using isoflurane and sacrificed by severing the spinal cord. Hippocampi were dissected from 3-6 ARL13B-EGFP^tg^ mice (ages P0-P1, sex undetermined, background strain C57BL/6J) and digested in ~20 units of papain (LK003176, Worthington) and ~200 units of deoxyribonuclease (LK003172, Worthington) dissolved to basal medium eagle solution (B1522, Sigma) at 37 °C for 25 min^40^. Tissues were washed with BME and triturated to release cells. Cells were centrifuged at 300 × g for 4 min and plated on polylysine-coated glass coverslip at 1-4 × 10^5^ density in basal medium eagle solution containing B27 supplement (17504044, Gibco), N-2 supplement (17502048, Gibco), 0.5% penicillin/streptomycin (15140148, Gibco), 5% fetal bovine serum, 5% horse serum (260500,Gibco), and GlutaMax (350500, Gibco). Hippocampal neurons were cultured for 2 to 7 days prior to conducting electrophysiology experiments. Inner medullary collecting duct (IMCD) cells stably expressing ARL13B-EGFP cilia reporters were cultured in F12/DMEM media (Sigma) with 10% fetal bovine serum (Sigma) and 50 I.U./mL penicillin-streptomycin (30-2300, ATCC) antibiotic.

### Electrophysiology

All research chemicals used in the electrophysiology experiments were purchased from Millipore-Sigma. Single-channel recordings were recorded from primary cilia and GUV membranes. All GUV patch electrodes were made using borosilicate glass electrodes and were fire polished to resistances greater than 5 - 10 MΩ and primary cilia patch electrodes were polished to a resistance greater than 15 MΩ. Renal primary cilia PKD2 currents were recorded from mIMCD-3 (ATCC, Catalog no. CRL-2123) cells expressing the cilia reporter ARL13B-EGFP, as previously described^48, 61-64^. Primary cilia PKD2L1 currents were recorded from isolated neonatal hippocampal neurons from ARL13B-EGFP^tg^ using previously described procedure^40^. Recording solutions for mammalian culture consisted of symmetrical recording solutions with 150 mM KCl, 10 mM HEPES, and 300 mM glucose, unless the charge carrier was changed when mentioned. Recordings were collected in voltage-clamp with voltage-steps from -100 mV to +100 mV and a holding potential of -40 mV with ClampEx v.11.2.0.59 (MolecularDevices, San Jose, CA USA) using a Axopatch 200B amplifier. Recordings were digitized with the Digidata 1550B (MolecularDevices) at 25kHz and low pass filtered at 5kHz. Recordings were analyzed with ClampFit v11.2.0.59 (Molecular Devices, San Jose, CA USA) and IGOR Pro 8.04 (Wavemetrics, Portland, OR USA). As a predetermined criteria, data was excluded from analysis when seal resistance fell below 5 MΩ due insufficient voltage control of the patched membrane. Conductance was determined by determining the slope of the current-voltage relationship. Probability of open time was calculated by measuring the time at which a channel spends in an open confirmation divided by the total time in the voltage step.

### Materials availability statement

All CFE and mammalian cell expression constructs used in this study are available without restriction upon written request to the corresponding author.

## Data availability statement

Data reported in this paper is deposited and available without restriction at the NU library ARCH (https://doi.org/10.21985/n2-4hs8-6j16).

## ACKNOWLEDGEMENTS

We acknowledge members of the Kamat and DeCaen labs for their constructive comments during the drafting of this manuscript. We thank Dr Alfred George for the use of his lipid electroformation equipment used to generate SUVs and GUVs. M.L was supported by Northwestern University’s (NU) molecular biophysics training grant (T32 GM008382) and the National Institute of Diabetes and Digestive and Kidney Diseases (NIDDK) of the National Institutes of Health kidney, urologic and hematologic (KUH) disease training grant (U2CDK129917). O.E.P. was supported by the Ruth L. Kirschstein National Research Service Award (NRSA) individual postdoctoral fellowship (F32DK137477-01A1) and NU KUH training grants (U2CDK129917 and TL1DK132769); P.G.D. was supported by the NIH NIDDK grants R01 DK123463-01 and R01 DK131118-01.

## FIGURE LEGENDS

**Figure 1— figure supplement 1. Quantification, identification and assembly of cell-free synthesized polycystin channels. A**). *Top*, Fluorescence (488γ) standard curve determined with a recombinant GFP tagged protein fit with a linear regression. PURExpress synthesized PKD2L1-GFP and PKD2-GFP were measured after three hours of expression at 37°C. (GFP standard curve N=5, PKD2L1 and PKD2 N=3 replicates). *Bottom*, average protomer and tetramer protein production from the PURExpress reaction. Error bars represent SEM. **B**) Representative sequence coverage of PKD2L1-GFP (top) and PKD2-GFP (bottom) from tandem mass spectrometry spectra. **C**) Mass spectrometry outputs identifying polycystin proteins. **D, E**) Fluorescence-detection size-exclusion chromatography (FSEC) of polycystin proteins derived from recombinant and CFE sources. Recombinant human PKD2-GFP and PKD2L1-GFP protein was obtained from lysates of 0.5×10^6^ HEK cells stably expressing the channels. Purified *Aequorea Victoria* GFP His-tag protein was obtained from ThermoFisher Scientific. SUVs containing CFE derived polycystins were lysed using dodecyl β-D-maltoside (DDM) prior to FSEC analysis (see methods).

**Figure 2— figure supplement 1. The SNAP-fluorescence approach to assess polycystin protein orientation in GUV**. Schematic of PKD2L1-SNAP incorporated into GUVs, followed by SNAP-staining with cell permeable (Cell488), and cell impermeable (Surface647) SNAP-Tag marker.

**Figure 3—figure supplement 1. Synthetic polycystin channels exhibit full and sub-conductive states in GUVs. A, E)** Single channel current amplitudes measured from individual GUVs (open circles). GUVs with open channel events measure at four or more potentials in inward and outward direction were fit to a linear equation to estimate their conductance. **B, F)** Resulting violin plots of the sub (*SC*, blue) and full (*FC*, black) polycystin conductance as estimated from individual GUV recordings (N = 5-7 GUVs). **C, G)** Unitary single channel currents measured from GUVs expressing polycystins held at 100 mV. Examples on the top demonstrated channels transitioning from closed (*C*) to the full conductance current levels, whereas examples below are measured from channels shifting between sub- and fully conductive current levels. **D, H**) Resulting histogram analysis of the corresponding single channel currents.

**Figure 4—figure supplement 1. Native polycystin channels measured from primary cilia membranes exhibit full and subconductance states. A, B**) *Top*, images of voltage clamped primary cilia from mouse hippocampal neurons and inner medullary kidney collecting duct cells (IMCD) harvested from transgenic mice expressing a fluorescent cilia reporter (ARL13B-EGFP^tg^)^65^. Previous work has genetically identified PKD2L1 and PKD2 as essential ion channel subunit in the primary cilia of the renal collecting duct cells and hippocampal neurons^40, 48^. *Bottom*, average single-channel current amplitudes recorded from primary cilia using K^+^ in the recording electrode solution. Conductance (γ) estimated by fitting the average single channel currents to a linear equation. Error indicates SEM (N=6 cilia).

## REFERENCES

1. Hille, B., Ionic channels in excitable membranes. Current problems and biophysical approaches. Biophysical journal 1978, 22 (2), 283–294.

2. Lee, A.; Fakler, B.; Kaczmarek, L. K.; Isom, L. L., More than a pore: ion channel signaling complexes. J Neurosci 2014, 34 (46), 15159–15169.

3. Yang, Y.; Wang, Z.; Gan, C.; Klausen, L. H.; Bonne, R.; Kong, G.; Luo, D.; Meert, M.; Zhu, C.; Sun, G.; Guo, J.; Ma, Y.; Bjerg, J. T.; Manca, J.; Xu, M.; Nielsen, L. P.; Dong, M., Long-distance electron transfer in a filamentous Gram-positive bacterium. Nat Commun 2021, 12 (1), 1709.

4. Prindle, A.; Liu, J.; Asally, M.; Ly, S.; Garcia-Ojalvo, J.; Suel, G. M., Ion channels enable electrical communication in bacterial communities. Nature 2015, 527 (7576), 59–63.

5. Goaillard, J. M.; Marder, E., Ion Channel Degeneracy, Variability, and Covariation in Neuron and Circuit Resilience. Annu Rev Neurosci 2021, 44, 335–357.

6. Stutzmann, G. E.; Mattson, M. P., Endoplasmic reticulum Ca(2+) handling in excitable cells in health and disease. Pharmacol Rev 2011, 63 (3), 700–27.

7. Smith, J. J.; Aitchison, J. D., Peroxisomes take shape. Nat Rev Mol Cell Biol 2013, 14 (12), 803–17.

8. Li, P.; Gu, M.; Xu, H., Lysosomal Ion Channels as Decoders of Cellular Signals. Trends Biochem Sci 2019, 44 (2), 110–124.

9. Yu, F. H.; Yarov-Yarovoy, V.; Gutman, G. A.; Catterall, W. A., Overview of molecular relationships in the voltage-gated ion channel superfamily. Pharmacol Rev 2005, 57 (4), 387–95.

10. George, A. L., Jr., Lessons learned from genetic testing for channelopathies. Lancet Neurol 2014, 13 (11), 1068–1070.

11. George, A. L., Jr., Recent genetic discoveries implicating ion channels in human cardiovascular diseases. Curr Opin Pharmacol 2014, 15, 47–52.

12. Clare, J. J., Targeting voltage-gated sodium channels for pain therapy. Expert Opin Investig Drugs 2010, 19 (1), 45–62.

13. Lieve, K. V.; Wilde, A. A., Inherited ion channel diseases: a brief review. Europace 2015, 17 Suppl 2, ii1–6.

14. Kass, R. S., The channelopathies: novel insights into molecular and genetic mechanisms of human disease. J Clin Invest 2005, 115 (8), 1986–9.

15. Overington, J. P.; Al-Lazikani, B.; Hopkins, A. L., How many drug targets are there? Nat Rev Drug Discov 2006, 5 (12), 993–6.

16. Santos, R.; Ursu, O.; Gaulton, A.; Bento, A. P.; Donadi, R. S.; Bologa, C. G.; Karlsson, A.; Al-Lazikani, B.; Hersey, A.; Oprea, T. I.; Overington, J. P., A comprehensive map of molecular drug targets. Nat Rev Drug Discov 2017, 16 (1), 19–34.

17. Kaczorowski, G. J.; McManus, O. B.; Priest, B. T.; Garcia, M. L., Ion channels as drug targets: the next GPCRs. J Gen Physiol 2008, 131 (5), 399–405.

18. Oprea, T. I., Exploring the dark genome: implications for precision medicine. Mamm Genome 2019, 30 (7-8), 192–200.

19. McGivern, J. G.; Ding, M., Ion Channels and Relevant Drug Screening Approaches. SLAS Discov 2020, 25 (5), 413–419.

20. Neher, E.; Sakmann, B., Single-channel currents recorded from membrane of denervated frog muscle fibres. Nature 1976, 260 (5554), 799–802.

21. Neher, E., Ion channels for communication between and within cells. Science 1992, 256 (5056), 498–502.

22. Hamill, O. P.; Marty, A.; Neher, E.; Sakmann, B.; Sigworth, F. J., Improved patch-clamp techniques for high-resolution current recording from cells and cell-free membrane patches. Pflugers Arch 1981, 391 (2), 85–100.

23. Morera, F. J.; Vargas, G.; Gonzalez, C.; Rosenmann, E.; Latorre, R., Ion-channel reconstitution. Methods Mol Biol 2007, 400, 571–85.

24. Leptihn, S.; Thompson, J. R.; Ellory, J. C.; Tucker, S. J.; Wallace, M. I., In vitro reconstitution of eukaryotic ion channels using droplet interface bilayers. J Am Chem Soc 2011, 133 (24), 9370–5.

25. Varghese, A.; Tenbroek, E. M.; Coles, J., Jr.; Sigg, D. C., Endogenous channels in HEK cells and potential roles in HCN ionic current measurements. Prog Biophys Mol Biol 2006, 90 (1-3), 26–37.

26. Pablo, J. L.; DeCaen, P. G.; Clapham, D. E., Progress in ciliary ion channel physiology. J Gen Physiol 2017, 149 (1), 37–47.

27. Shimizu, Y.; Ueda, T., PURE technology. Methods Mol Biol 2010, 607, 11–21.

28. Kuruma, Y.; Nishiyama, K.; Shimizu, Y.; Müller, M.; Ueda, T., Development of a minimal cell-free translation system for the synthesis of presecretory and integral membrane proteins. Biotechnol Prog 2005, 21 (4), 1243–51.

29. Jacobs, M. L.; Kamat, N. P., Cell-Free Membrane Protein Expression into Hybrid Lipid/Polymer Vesicles. Methods Mol Biol 2022, 2433, 257–271.

30. Shen, P. S.; Yang, X.; DeCaen, P. G.; Liu, X.; Bulkley, D.; Clapham, D. E.; Cao, E., The Structure of the Polycystic Kidney Disease Channel PKD2 in Lipid Nanodiscs. Cell 2016, 167 (3), 763–773 e11.

31. Grieben, M.; Pike, A. C.; Shintre, C. A.; Venturi, E.; El-Ajouz, S.; Tessitore, A.; Shrestha, L.; Mukhopadhyay, S.; Mahajan, P.; Chalk, R.; Burgess-Brown, N. A.; Sitsapesan, R.; Huiskonen, J. T.; Carpenter, E. P., Structure of the polycystic kidney disease TRP channel Polycystin-2 (PC2). Nat Struct Mol Biol 2017, 24 (2), 114–122.

32. Wang, Q.; Corey, R. A.; Hedger, G.; Aryal, P.; Grieben, M.; Nasrallah, C.; Baronina, A.; Pike, A. C. W.; Shi, J.; Carpenter, E. P.; Sansom, M. S. P., Lipid Interactions of a Ciliary Membrane TRP Channel: Simulation and Structural Studies of Polycystin-2. Structure 2020, 28 (2), 169–184 e5.

33. Wilkes, M.; Madej, M. G.; Kreuter, L.; Rhinow, D.; Heinz, V.; De Sanctis, S.; Ruppel, S.; Richter, R. M.; Joos, F.; Grieben, M.; Pike, A. C.; Huiskonen, J. T.; Carpenter, E. P.; Kuhlbrandt, W.; Witzgall, R.; Ziegler, C., Molecular insights into lipid-assisted Ca(2+) regulation of the TRP channel Polycystin-2. Nat Struct Mol Biol 2017, 24 (2), 123–130.

34. Hulse, R. E.; Li, Z.; Huang, R. K.; Zhang, J.; Clapham, D. E., Cryo-EM structure of the polycystin 2-l1 ion channel. Elife 2018, 7.

35. Su, Q.; Hu, F.; Liu, Y.; Ge, X.; Mei, C.; Yu, S.; Shen, A.; Zhou, Q.; Yan, C.; Lei, J.; Zhang, Y.; Liu, X.; Wang, T., Cryo-EM structure of the polycystic kidney disease-like channel PKD2L1. Nat Commun 2018, 9 (1), 1192.

36. Esarte Palomero, O.; Larmore, M.; DeCaen, P. G., Polycystin Channel Complexes. Annu Rev Physiol 2023, 85, 425–448.

37. Wu, G.; D’Agati, V.; Cai, Y.; Markowitz, G.; Park, J. H.; Reynolds, D. M.; Maeda, Y.; Le, T. C.; Hou, H., Jr.; Kucherlapati, R.; Edelmann, W.; Somlo, S., Somatic inactivation of Pkd2 results in polycystic kidney disease. Cell 1998, 93 (2), 177–88.

38. Gao, Z.; Ruden, D. M.; Lu, X., PKD2 cation channel is required for directional sperm movement and male fertility. Curr Biol 2003, 13 (24), 2175–8.

39. Tanaka, Y.; Morozumi, A.; Hirokawa, N., Nodal flow transfers polycystin to determine mouse left-right asymmetry. Dev Cell 2023, 58 (16), 1447–1461 e6.

40. Vien, T. N.; Ta, M. C.; Kimura, L. F.; Onay, T.; DeCaen, P. G., Primary cilia TRP channel regulates hippocampal excitability. Proc Natl Acad Sci U S A 2023, 120 (22), e2219686120.

41. Yao, G.; Luo, C.; Harvey, M.; Wu, M.; Schreiber, T. H.; Du, Y.; Basora, N.; Su, X.; Contreras, D.; Zhou, J., Disruption of polycystin-L causes hippocampal and thalamocortical hyperexcitability. Hum Mol Genet 2016, 25 (3), 448–58.

42. Vo, C. V.; Mikutis, G.; Bode, J. W., SnAP reagents for the transformation of aldehydes into substituted thiomorpholines--an alternative to cross-coupling with saturated heterocycles. Angew Chem Int Ed Engl 2013, 52 (6), 1705–8.

43. Müller-Lucks, A.; Bock, S.; Wu, B.; Beitz, E., Fluorescent in situ folding control for rapid optimization of cell-free membrane protein synthesis. PLoS One 2012, 7 (7), e42186.

44. Klammt, C.; Schwarz, D.; Löhr, F.; Schneider, B.; Dötsch, V.; Bernhard, F., Cell-free expression as an emerging technique for the large scale production of integral membrane protein. Febs j 2006, 273 (18), 4141–53.

45. Boban, Z.; Mardesic, I.; Subczynski, W. K.; Raguz, M., Giant Unilamellar Vesicle Electroformation: What to Use, What to Avoid, and How to Quantify the Results. Membranes (Basel) 2021, 11 (11).

46. DeCaen, P. G.; Liu, X.; Abiria, S.; Clapham, D. E., Atypical calcium regulation of the PKD2-L1 polycystin ion channel. Elife 2016, 5.

47. Ng, L. C. T.; Vien, T. N.; Yarov-Yarovoy, V.; DeCaen, P. G., Opening TRPP2 (PKD2L1) requires the transfer of gating charges. Proc Natl Acad Sci U S A 2019, 116 (31), 15540–15549.

48. Kleene, S. J.; Kleene, N. K., The native TRPP2-dependent channel of murine renal primary cilia. Am J Physiol Renal Physiol 2017, 312 (1), F96-F108.

49. Liu, X.; Vien, T.; Duan, J.; Sheu, S. H.; DeCaen, P. G.; Clapham, D. E., Polycystin-2 is an essential ion channel subunit in the primary cilium of the renal collecting duct epithelium. Elife 2018, 7.

50. Cai, Y.; Maeda, Y.; Cedzich, A.; Torres, V. E.; Wu, G.; Hayashi, T.; Mochizuki, T.; Park, J. H.; Witzgall, R.; Somlo, S., Identification and characterization of polycystin-2, the PKD2 gene product. J Biol Chem 1999, 274 (40), 28557–65.

51. Newby, L. J.; Streets, A. J.; Zhao, Y.; Harris, P. C.; Ward, C. J.; Ong, A. C., Identification, characterization, and localization of a novel kidney polycystin-1-polycystin-2 complex. J Biol Chem 2002, 277 (23), 20763–73.

52. Nakatsu, F., A Phosphoinositide Code for Primary Cilia. Dev Cell 2015, 34 (4), 379–80.

53. Noireaux, V.; Liu, A. P., The New Age of Cell-Free Biology. Annu Rev Biomed Eng 2020, 22, 51–77.

54. Gopfrich, K.; Haller, B.; Staufer, O.; Dreher, Y.; Mersdorf, U.; Platzman, I.; Spatz, J. P., One-Pot Assembly of Complex Giant Unilamellar Vesicle-Based Synthetic Cells. ACS Synth Biol 2019, 8 (5), 937–947.

55. Zhou, X.; Belavek, K. J.; Miller, E. W., Origins of Ca(2+) Imaging with Fluorescent Indicators. Biochemistry 2021, 60 (46), 3547–3554.

56. Yu, H. B.; Li, M.; Wang, W. P.; Wang, X. L., High throughput screening technologies for ion channels. Acta Pharmacol Sin 2016, 37 (1), 34–43.

57. Li, K.; Jiang, Q.; Bai, X.; Yang, Y. F.; Ruan, M. Y.; Cai, S. Q., Tetrameric Assembly of K(+) Channels Requires ER-Located Chaperone Proteins. Mol Cell 2017, 65 (1), 52–65.

58. Chen, Z.; Mondal, A.; Abderemane-Ali, F.; Jang, S.; Niranjan, S.; Montano, J. L.; Zaro, B. W.; Minor, D. L., Jr., EMC chaperone-Ca(V) structure reveals an ion channel assembly intermediate. Nature 2023, 619 (7969), 410–419.

59. Bai, X.; Li, K.; Yao, L.; Kang, X. L.; Cai, S. Q., A forward genetic screen identifies chaperone CNX-1 as a conserved biogenesis regulator of ERG K(+) channels. J Gen Physiol 2018, 150 (8), 1189–1201.

60. Vien, T. N.; Wang, J.; Ng, L. C. T.; Cao, E.; DeCaen, P. G., Molecular dysregulation of ciliary polycystin-2 channels caused by variants in the TOP domain. Proc Natl Acad Sci U S A 2020, 117 (19), 10329–10338.

61. Kleene, N. K.; Kleene, S. J., A method for measuring electrical signals in a primary cilium. Cilia 2012, 1.

62. DeCaen, P. G.; Delling, M.; Vien, T. N.; Clapham, D. E., Direct recording and molecular identification of the calcium channel of primary cilia. Nature 2013, 504 (7479), 315–8.

63. Liu, X.; Vien, T.; Duan, J.; Sheu, S.-H.; DeCaen, P. G.; Clapham, D. E., Polycystin-2 is an essential ion channel subunit in the primary cilium of the renal collecting duct epithelium. Elife 2017.

64. Vien, T. N.; Wang, J.; Ng, L. C. T.; Cao, E.; DeCaen, P. G., Molecular dysregulation of ciliary polycystin-2 channels caused by variants in the TOP domain. Proc Natl Acad Sci U S A 2020.

65. Delling, M.; DeCaen, P. G.; Doerner, J. F.; Febvay, S.; Clapham, D. E., Primary cilia are specialized calcium signalling organelles. Nature 2013, 504 (7479), 311–4.

