## Supplemental Figures for "A synthetic method to assay polycystin channel biophysics"

A

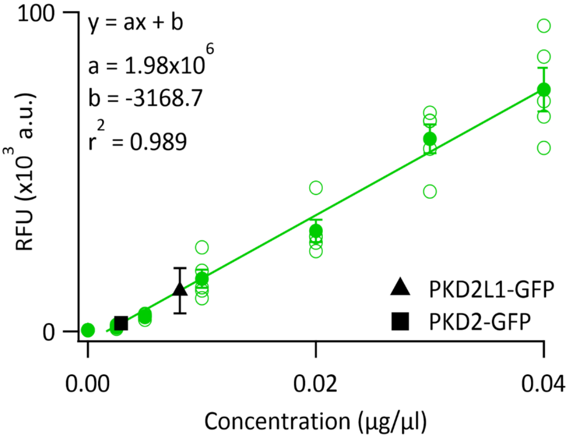


| | Average protomer [ng/ul] | Average tetramer [ng/ul] |
|---|---|---|
| PKD2L1-GFP | 8.0 ± 3.6 | 2.0 ± 0.9 |
| PKD2-GFP | 3.1 ± 2 | 0.8 ± 0.07 |

B

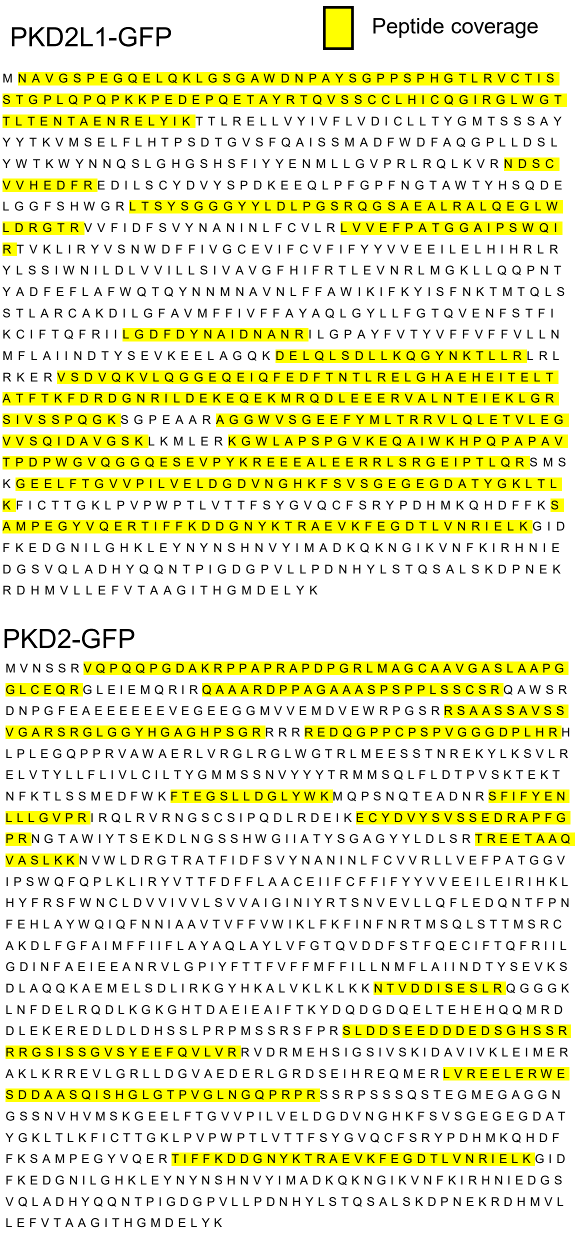


C

| | Protein ID | Percent coverage | Exclusive unique peptides |
|---|---|---|---|
| PKD2L1-GFP | 100% | 46% | 22 |
| PKD2-GFP | 100% | 25% | 63 |

D

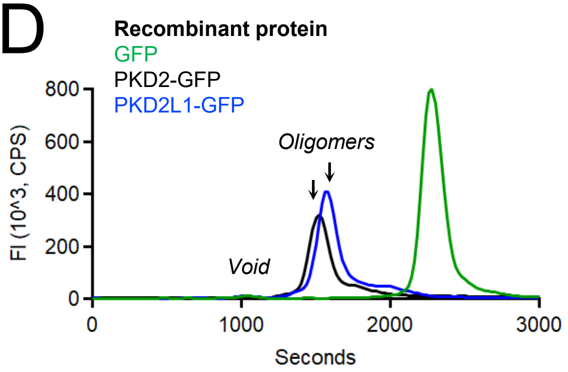


E

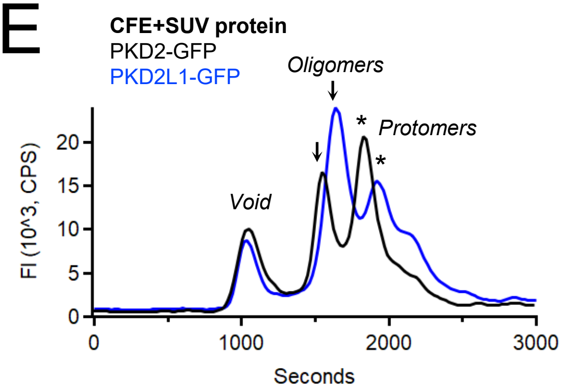


Figure 1—Figure Supplement 1

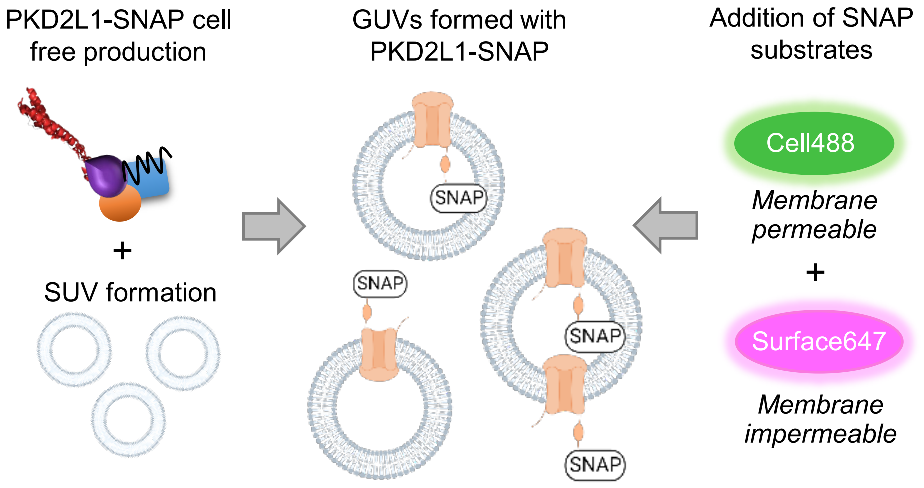


Figure 2– Figure supplement 1

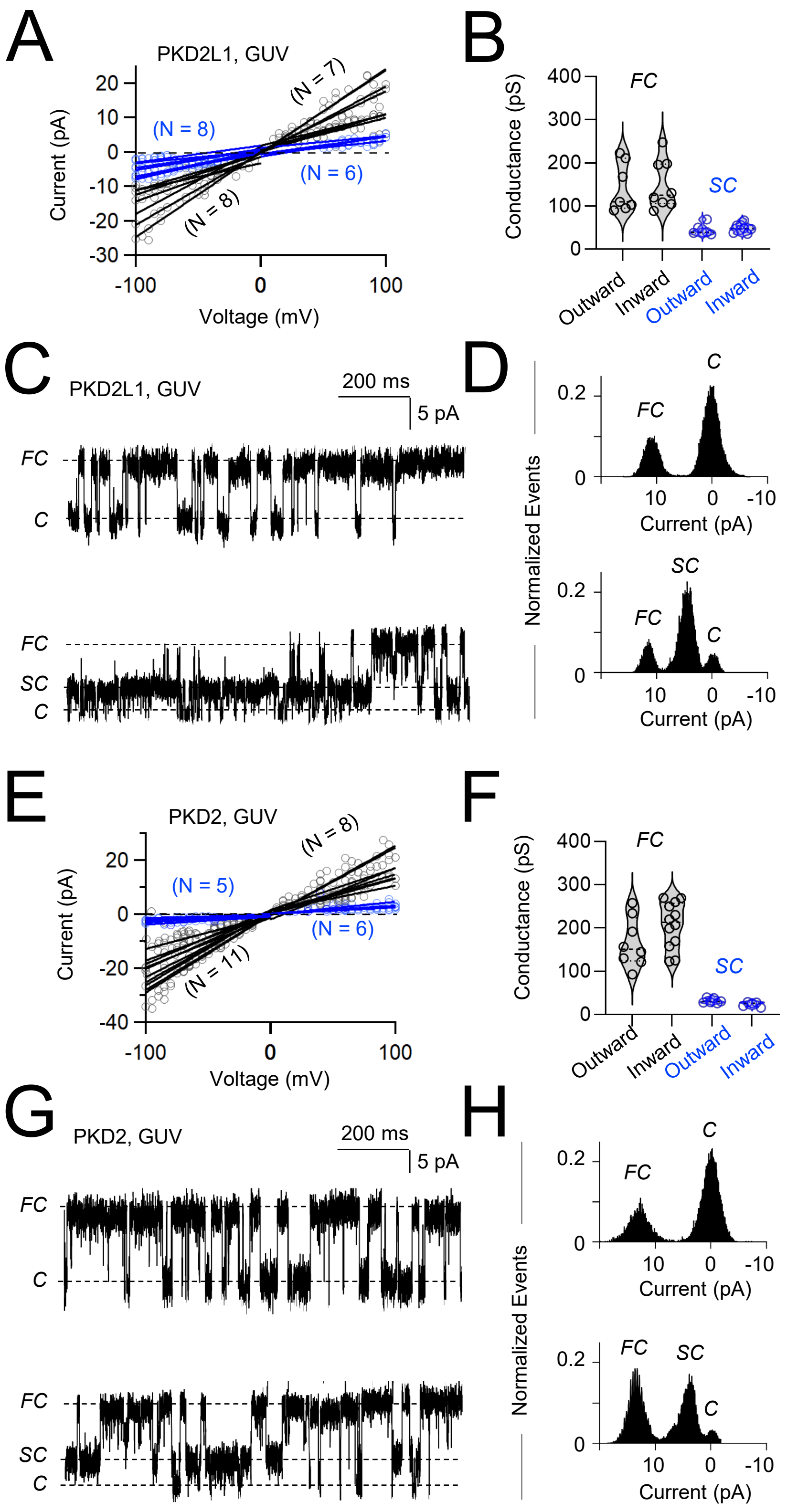


Figure 3– Figure Supplement 1

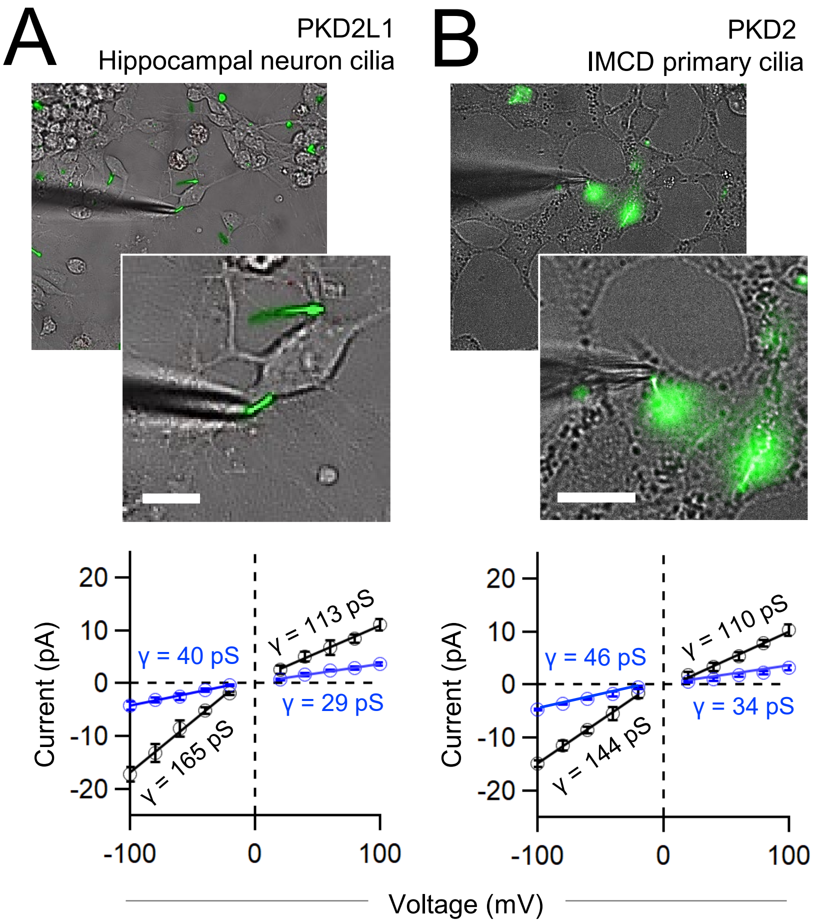


Figure 4– Figure supplement 1
